# A scalable platform for efficient CRISPR-Cas9 chemical-genetic screens of DNA damage-inducing compounds

**DOI:** 10.1101/2023.06.16.545157

**Authors:** Kevin Lin, Ya-Chu Chang, Maximilian Billmann, Henry N. Ward, Khoi Le, Arshia Hassan, Urvi Bhojoo, Katherine Chan, Michael Costanzo, Jason Moffat, Charles Boone, Anja-Katrin Bielinsky, Chad L. Myers

## Abstract

Current approaches to define chemical-genetic interactions (CGIs) in human cell lines are resource-intensive. We designed a scalable chemical-genetic screen platform by generating a DNA damage response (DDR)-focused custom sgRNA library. We performed five proof-of-principle compound screens and found that the compounds’ known modes-of-action (MoA) were enriched among the compounds’ CGIs. These scalable screens recapitulated expected CGIs at a comparable signal-to-noise ratio (SNR) relative to genome-wide screens. Furthermore, time-resolved CGIs, captured by sequencing screens at various time points, suggested an unexpected, late time point interstrand-crosslinking (ICL) repair pathway response to camptothecin-induced DNA damage. Our approach can facilitate screening compounds at scale and produce biologically informative CGI profiles.

## Introduction

Screening chemical compounds against a collection of defined gene knockouts can identify mutants that sensitize or suppress a compound’s phenotypic effect^1^. This approach, known as chemical-genetic interaction (CGI) profiling, has relevant clinical applications for discovering novel genetic vulnerabilities or resistance mechanisms in the context of existing targeted therapies, particularly in cancer^2^.

Many chemical-genetic screens have been performed in *S. cerevisiae*, a model organism amenable to facile genetic manipulation^3^. *S. cerevisiae* gene deletion libraries can be constructed such that each strain harbors a specific gene knockout, and collections of yeast mutant strains can be easily screened against chemical compound libraries in a high-throughput manner^4^. A phenotypic output, such as cell fitness, can be quantified from these chemical-genetic screens to determine if a certain gene knockout confers sensitivity (negative CGI) or resistance (positive CGI) to a compound. This unbiased approach to chemical-genetic screens in *S. cerevisiae*, which produced chemical-genetic fingerprint profiles using a small subset of the genome-wide deletion library, has led to mode-of-action (MoA) predictions for thousands of compounds^5^.

Similar chemical-genetic screens have been developed in human cell line models, with early approaches adopting a RNA interference (RNAi) knockdown strategy with short hairpin RNA (shRNA) libraries^6^. More recently, the advent of clustered regularly interspaced short palindromic repeats (CRISPR)/CRISPR-associated protein 9 (Cas9) editing technology allowed for facile construction of gene knockouts^7, 8^. Pooled lentiviral CRISPR-Cas9 screening using single guide RNA (sgRNA) libraries enable interrogation of gene knockout phenotypes on a genome-wide scale in human cell lines^9, 10^.

Pooled chemical-genetic CRISPR screens have been adopted in human cell lines as an analogous method to chemical-genetic screening in *S. cerevisiae*^11^. Several large-scale chemical screens have been performed in human cell lines, including efforts to map the DNA damage response network^12, 13^ or to characterize the ubiquitin-proteasome system^14, 15^. However, there is currently an unmet need for a scalable and generalizable approach towards a low-cost, high-throughput chemical screening method in human cell lines. Moreover, resolution on certain technical parameters for CRISPR chemical screens, such as compound dosage, intermediate time points, and library representation, have not been thoroughly investigated.

Screening well-characterized genotoxins can provide insight into the applicability of this novel approach. Genotoxins cause DNA lesions, which, if not repaired correctly, lead to mutations or genomic aberrations that threaten cell viability^16^. The DNA damage response (DDR), a network of damage signaling pathways and DNA repair pathways working in concert, promotes the sensing and repair of DNA lesions and prevents genomic instability. While the general mechanisms of this network have been well described, much of this network complexity has not been elucidated. Specifically, CGI profiling of genotoxins against DDR genes may provide better understanding of synthetic lethal interactions that can be exploited for combination therapies, or mechanisms of resistance to chemotherapies.

Here, we propose a chemical CRISPR screening platform that takes advantage of a compressed, DDR-focused library. The reduction in costs, particularly for cell culture reagents and next-generation sequencing, allows for a scalable approach to screening a large number of compounds^17^. We performed 5 proof-of-principle screens against genotoxins or compounds that interact with the DDR network. Our screens recapitulated expected CGIs at a similar signal-to-noise ratio (SNR) compared to genome-wide screens and showed that CGIs are enriched in genes related to the characterized mechanism of action of a particular compound. Notably, our scalable screening approach also discovered previously unreported CGIs. Moreover, intermediate time point CGI data revealed novel time-resolved dependency of DNA repair pathways.

## Results

### Development of a targeted library for scalable CRISPR screens

Previous work in *S. cerevisiae* demonstrated that mutants covering a small subset of the genome were able to generate chemical-genetic fingerprints representative of a compound’s MoA^5^. With the long-term goal of establishing an analogous, highly scalable chemical genetic screening platform for human cells, we developed a proof-of-concept small, custom sgRNA lenti-library (hereafter referred to as the “targeted library”) for efficient chemical genetic screens. Our targeted library was designed to target four general categories of genes (Fig. 1a): well-characterized DNA damage response genes (n = 349), genes that captured the greatest variance across published CRISPR screens (n = 100), genes that captured subtle fitness defects in CRISPR screen data (n = 216), and genes that have a high degree of genetic interactions, or frequent interactors (n = 463) (see *Methods* for details on each category). Overall, the targeted library contained 3,033 sgRNAs targeting 1,011 genes (3 independent sgRNAs/gene). The sgRNA library was optimally selected from a pool of guides from the genome-wide Toronto KnockOut version 3.0 (TKOv3) library, which contains ∼71,000 guides targeting ∼18,000 genes (4 independent sgRNAs/gene)^18^. Selection of library genes was based on previous screen data and quality metrics (see *Methods*). The small library size enabled each replicate screen to be conducted on a series of single 15-cm tissue culture plates while being sufficient for maintaining a robust 1000X representation of each sgRNA in this scalable experimental format, compared to the 250-400X representation commonly used in other published CRISPR screens^12^. Overall, we estimate that this library provides a >20-fold increase in the number of compounds or cell lines that can be screened for the same cost.

**Fig. 1.**
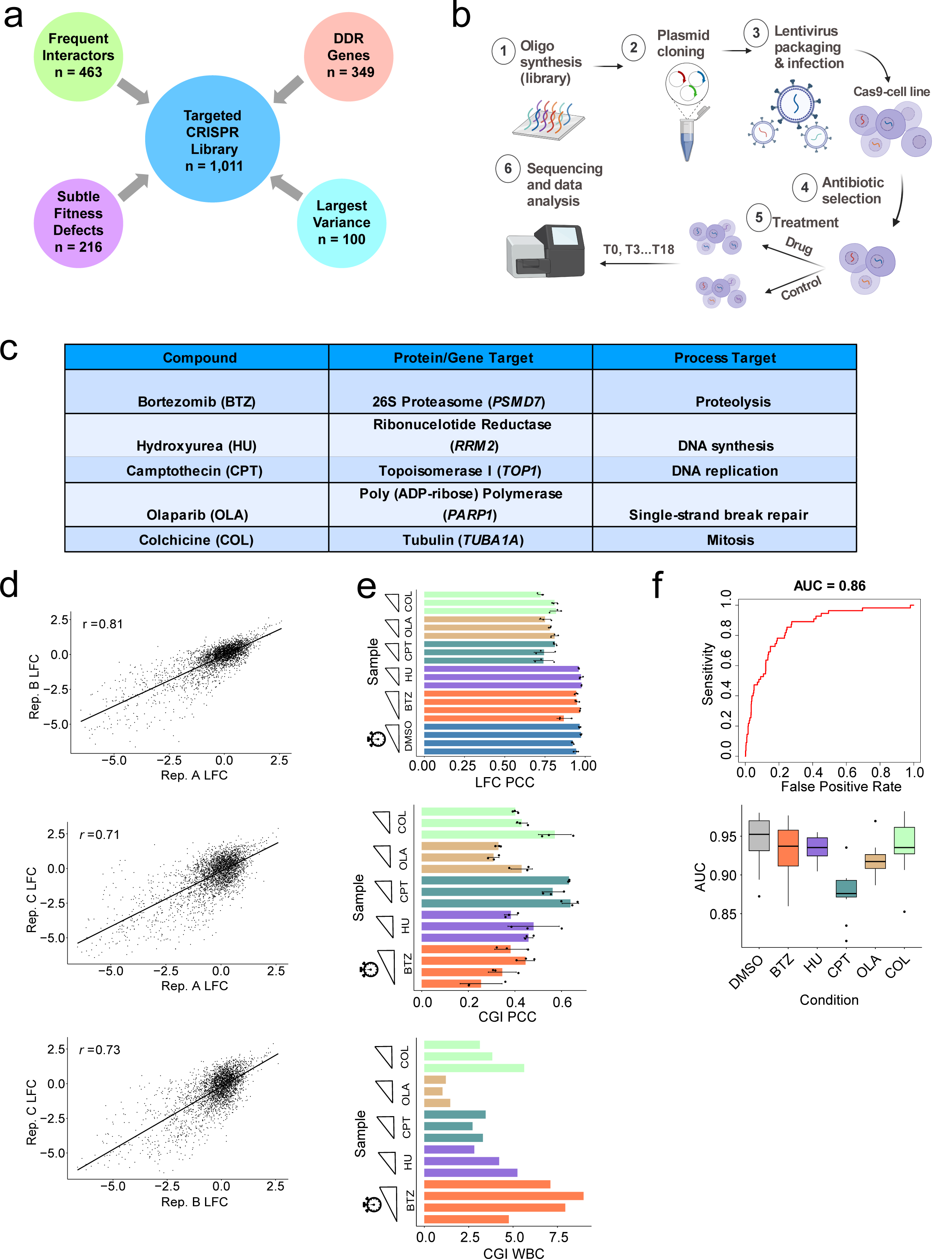
Scalable CRISPR-Cas9 chemical screen platform. **a** The categories of genes selected for the targeted sgRNA library. Number of genes (n) are indicated for each category. Genes can belong to more than one category. **b** Overview of scalable CRISPR-Cas9 chemical screen workflow. Schematic created with BioRender.com. **c** Table of 5 compounds used in proof-of-principle screens. For each compound, the protein target (corresponding gene italicized) and bioprocess target are listed. **d** Representative scatter plots of log_2_ fold change (LFC) values, or cell fitness, among 3 technical replicates of the camptothecin (CPT) T12 screen. Pearson’s correlation coefficient (*r*) is reported for each technical replicate pair (AB, AC, BC). Each point represents one sgRNA. **e** *Top*: Barplot of mean Pearson’s correlation coefficient (*r*) between vectors of LFC values for each replicate pair (AB, AC, BC). Points represent each *r* value, and standard deviation bars are overlaid. Sliding ramps represent increasing time points. *Middle*: Barplot of mean *r* between vectors of CGI scores for each replicate pair. *Bottom*: Barplot of within-vs-between context (WBC) correlation calculated on CGI scores. **f** *Top*: Receiver operating characteristic curve for discriminating essential vs. non-essential gene dropout for the T12 CPT screen. AUC: area under the curve. *Bottom*: Box-and-whisker plot displaying distribution of AUC values across control (DMSO) or compound screens. Each point represents the AUC value for each time point/screen replicate.

### Scalable CRISPR chemical screen workflow

To evaluate the utility of our targeted CRISPR library, we performed a set of 5 proof-of-principle pooled CRISPR-Cas9 chemical screens in the hTERT-immortalized RPE-1 *TP53* knockout cell line expressing a Flag-tagged Cas9 protein (Fig. 1b). Screens were conducted in a *TP53*-null background, as Cas9 cutting can induce a p53-mediated DNA damage response and cell cycle arrest, potentially masking the identification of essential gene dropouts in screens^19–21^. The 5 compounds (bortezomib - BTZ, hydroxyurea - HU, camptothecin - CPT, olaparib - OLA, and colchicine - COL) were selected due to: 1) the well-characterized protein and bioprocess targets of the compounds, and 2) the fact that genome-wide screen data for these compounds were either already publicly available or generated by us for comparison (Fig. 1c, Table 1). To determine the optimal dosage for these screens, we conducted a pilot screen with bortezomib at IC_50_ and IC_20_ (inhibitory concentrations determined after 3 days of exposure to compound by CellTiterGlo® cell viability assay). Dosing at IC_20_ (20% decrease in cell viability relative to vehicle treated only cells) vs. IC_50_ revealed similar CGIs (Fig. S1). While there was no definitive evidence to suggest using one dosage over the other, we reasoned that the IC_20_ dosage allowed a sufficiently large window to capture gene knockouts that either sensitize or suppress the compound’s effect on cell viability. In addition, the lower dosage ensured that enough cells could be passaged and collected throughout the length of the screen. The remaining proof-of-principle screens were performed with IC_20_ drug concentrations, as previously described^22^.

**Table 1:** Scalable vs. genome-wide screen parameters

|  | **Scalable Screens** | **Genome-Wide Screens** | **Genome-Wide Screens (Olivieri)** |
| --- | --- | --- | --- |
| **Compounds** | BTZ, HU, CPT, OLA, COL | CPT, OLA, COL | HU, CPT, OLA |
| **Dosage** | IC<sub>20</sub> | IC<sub>50</sub> | IC<sub>20</sub> * |
| **Timepoints** | T6, T9, T12, T15, T18 | T18 | T18 |
| **Coverage** | 1000X | 250X | 250-400X |
| **Replicates** | Triplicates | Triplicates | Duplicates |
| **Cell Line** | RPE-1 hTERT p53-/- | HAP1 | RPE-1 hTERT p53-/- |

**\*determined over 12 days instead of 3 days**

The NGS data was processed by an adapted version of a previously described computational tool for scoring CGIs^23^ (see *Methods* for details). Briefly, raw read counts were normalized by the sequencing depth of each sample. Guide fitness values were calculated as log_2_ fold changes (LFCs) at each time point relative to T0 for both untreated (DMSO) and treated (compound) conditions. Gene-level LFCs were calculated by averaging the guide-level LFCs across the 3 guides. CGI scores were quantified as a corrected differential LFC between treated and untreated conditions and scored for statistical significance using a moderated t-test^24^.

Screen quality was assessed through multiple metrics. First, we assessed how well the fitness defect data correlated between all pairwise replicates for a given screen (representative example for the T12 Camptothecin screen shown in Fig. 1d). Across all screens, the guide-level LFC data had high correlation among replicates (mean Pearson’s correlation coefficient (PCC), *r* = 0.8, Fig. 1e). Second, to better quantify the reproducibility of compound-specific effects, we assessed the correlation of the CGI values. The mean PCC for the CGI-scores was *r* = 0.24 (Fig. 1e). Given the sparsity of compound-specific effects from CRISPR screens, PCC is not a sensitive metric for assessing reproducibility of CGI scores^25^. Thus, we used the within-vs-between context correlation (WBC) score to quantify how similar CGI scores were among replicate screens relative to other screens^25^. This metric provided a more accurate representation of the reproducibility of the compound-specific effects, which was high for the majority of replicate screens (mean WBC: 4.85, WBC range: 1.03-8.97, Fig. 1e). In addition, essential genes dropped out as expected throughout the screen, as evaluated by a binary classification approach (mean area under the receiver operating characteristic curve metric, AUC-ROC = 0.9, Fig. 1f).

### Mapping chemical-genetic interactions with the targeted library

CGI scores are used to quantify whether library gene knockouts sensitize (negative CGI) or suppress/mask (positive CGI) a compound’s effect on cell viability. We defined significant CGI hits by using a CGI score (differential LFC) cutoff of greater than 0.7 (for positive CGIs) or less than −0.7 (for negative CGIs) and a false discovery rate (FDR) lower than 10% (Supplementary Table 1). Overall, 623 gene-compound interactions were identified across all time points, and ∼40% of the targeted library displayed interactions with at least one of the screened compounds for at least one time point. CGI scores correlate across time points for a given compound and are moderately correlated across the genotoxins (HU, CPT, OLA), reflecting their similar modes of action (Fig. S2).

Compound-specific CGIs recapitulate expected hits when considering the well-characterized bioprocess or pathway targets of these compounds, as shown on a global heatmap of the CGIs across all screens (Fig. 2a-f, S3-S4). Hydroxyurea, a potent ribonucleotide reductase inhibitor that depletes the dNTP pool and results in stalled replication forks, showed expected strong negative interactions with *RAD1*, *HUS1*, and *RAD17* (Fig. 2c). The Rad9-Hus1-Rad1 heterotrimeric complex (also known as the 9-1-1 complex) is a sliding clamp loaded by a complex containing Rad17 onto the DNA to sense sites of DNA damage and regulate checkpoint signaling pathways^26, 27^. *RAD9* was not included in the targeted library, and thus, does not appear as a negative CGI.

**Fig. 2.**
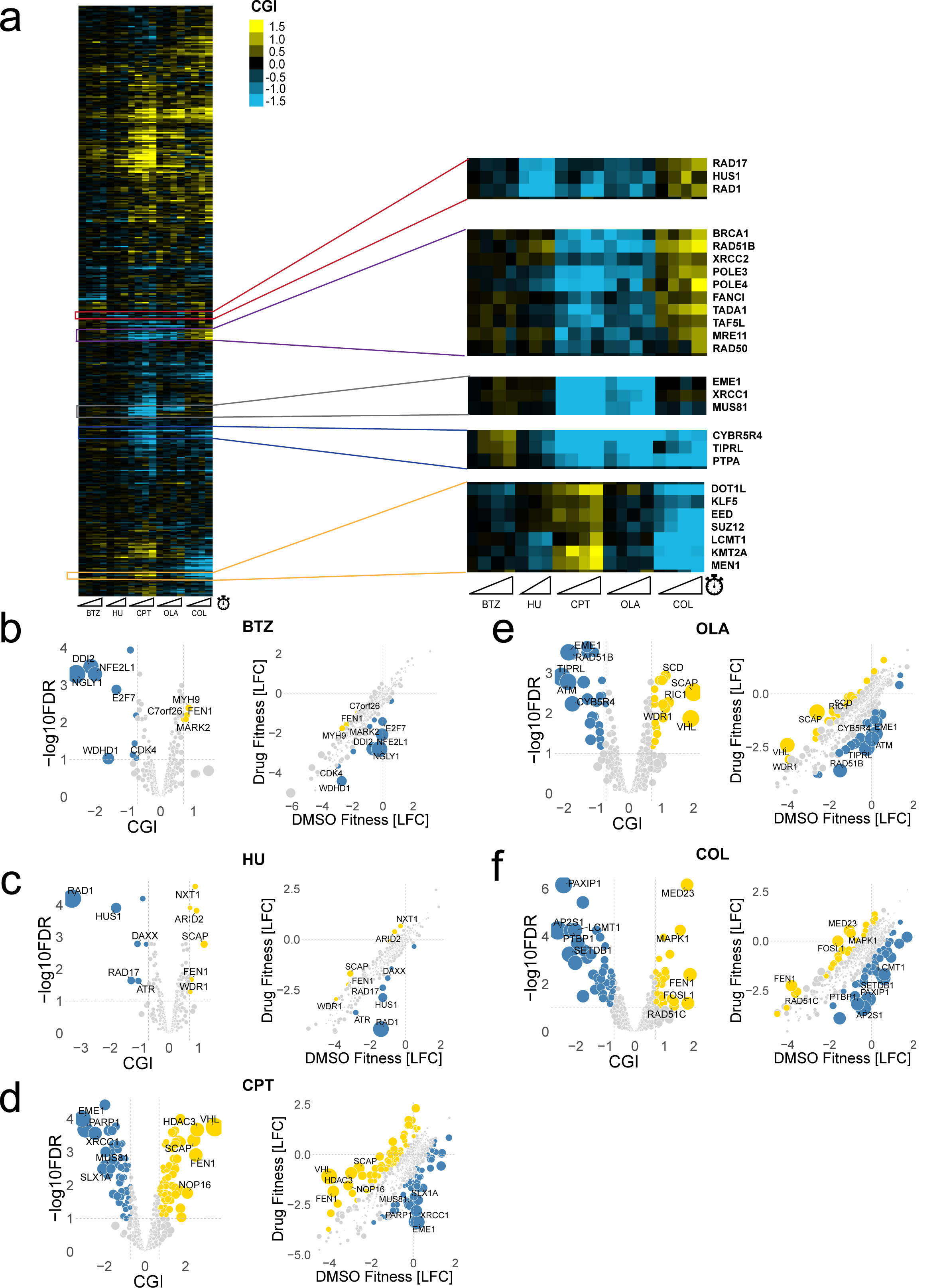
Proof-of-principle scalable chemical screens recapitulate expected chemical-genetic interactions. **a** Heatmap of CGI values across five compound screens. Heatmap uses average-linkage hierarchical clustering on gene side (rows), while each column represents a compound at specific time points (sliding ramps represent increasing time points). Blue pixels represent negative CGIs, and yellow pixels represent positive CGIs (saturated at |CGI| = 1.5). Representative clusters are enlarged. **b-f** *Left*: Volcano plot of T12 screen for each compound screen (BTZ = bortezomib, HU = hydroxyurea, CPT = camptothecin, OLA = olaparib, COL = colchicine). *Right*: Scatterplot of control (DMSO) fitness vs. compound fitness, as represented by LFC values. Negative and positive chemical-genetic interactions (CGIs) are indicated in blue and yellow, respectively. Each point represents a gene. False discovery rate (FDR) values were calculated using the Benjamini-Hochberg method. Cutoffs for significant CGIs (hits) were set at FDR = 0.1 and |CGI| > 0.7 (gray dashed lines). The top five negative and positive hits are labeled.

Camptothecin, a selective topoisomerase I inhibitor, and olaparib, a poly (ADP-ribose) polymerase (PARP) inhibitor, both covalently link and trap their respective enzyme targets to DNA, resulting in replication fork stalling and collapse, and DNA damage in the form of double-strand breaks^28, 29^. Both compounds showed strong negative interactions with members of the homologous recombination (HR) repair pathway, such as *BRCA1*, *RAD51B*, *XRCC1*, *XRCC2*, *MRE11*, *RAD50*, *EME1*, *MUS81*, and *RAD54L* (Fig. 2d-e). The MRE11-RAD50-NBS1 (MRN) complex, which has a role in sensing and repair of DNA damage, and Mus81-Eme1 endonuclease, which plays a role in processing stalled replication fork intermediates, may both be essential for DNA damage caused by camptothecin and olaparib^30, 31^. All genotoxins screened, including camptothecin, olaparib, and hydroxyurea, exhibit strong negative interactions with *CYB5R4*, which encodes an oxidoreductase and possible modulator of protein phosphatases, as well as with *TIPRL* and *PPP2R4*, which encode proteins that regulate the assembly and disassembly of protein phosphatase 2A (PP2A) complexes^12^ (Fig. 2a). These interactions were discovered in the genome-wide screens conducted by Olivieri et al. and recapitulated in our genotoxin scalable screens.

Colchicine, a beta-tubulin inhibitor that disrupts microtubule assembly and is often used to arrest cells in metaphase^32^, displayed strong positive interactions with the aforementioned HR genes (along with *RAD51D*, *XRCC3*), as well as with DNA replication genes such as *GINS4*, *MCM6*, *ORC2* (Fig. 2f). For cells arrested in mitosis, depletion of DNA repair and replication genes would not decrease the viability of these cells relative to untreated conditions. Interestingly, a group of chromatin remodeling genes (*DOT1L*, *EED*, *SUZ12*, *LCMT1*, *KMT2A*) displayed strong negative interactions with colchicine and strong positive interactions with camptothecin (Fig. 2a); these interactions have not been previously reported. EED and SUZ12 are members of the polycomb repressive complex 2 (PRC2), which is a histone methyltransferase that represses transcription through methylation of lysine 27 of histone H3 (H3K27)^33^. *KMT2A* encodes a histone methyltransferase that methylates H3K4, *DOT1L* encodes a lysine methyltransferase that methylates H3K79, and *LCMT1* encodes a leucine carboxyl methyltransferase that regulates PP2A methylation. Overall, the proof-of-principle compound screens recapitulated previously reported CGIs and revealed novel CGIs.

### Compound mode-of-action enriched in CGIs

We asked whether genes in the compounds’ known modes-of-action were enriched in their CGI hits. To perform this analysis, we first collated the protein targets of each compound (Fig. 3a). For each of these targets, we selected a GO:BP term that best describes the compound MoA, or the biological process targeted by the compound. Next, we investigated whether the genes in this targeted library annotated to these GO:BP terms were overrepresented in the significant hits for each compound screen. To interrogate whether the MoA was generally enriched across all compound screens, we combined compound-hit pairs across all compounds before conducting a statistical test and measuring fold enrichment. Across the set of all compounds’ CGI scores, MoA related genes were significantly enriched (fold enrichment = 1.52), with primarily negative CGIs driving this enrichment (Fig. 3b). Further subdividing CGIs into essential and non-essential genes (see *Methods*), revealed that this enrichment on negative interactions occurred regardless of essentiality status of the gene (Fig. 3c). Specifically, sgRNAs targeting essential genes related to the MoA tended to drop out more quickly in cells exposed to these compounds relative to the control (DMSO) condition.

**Fig. 3.**
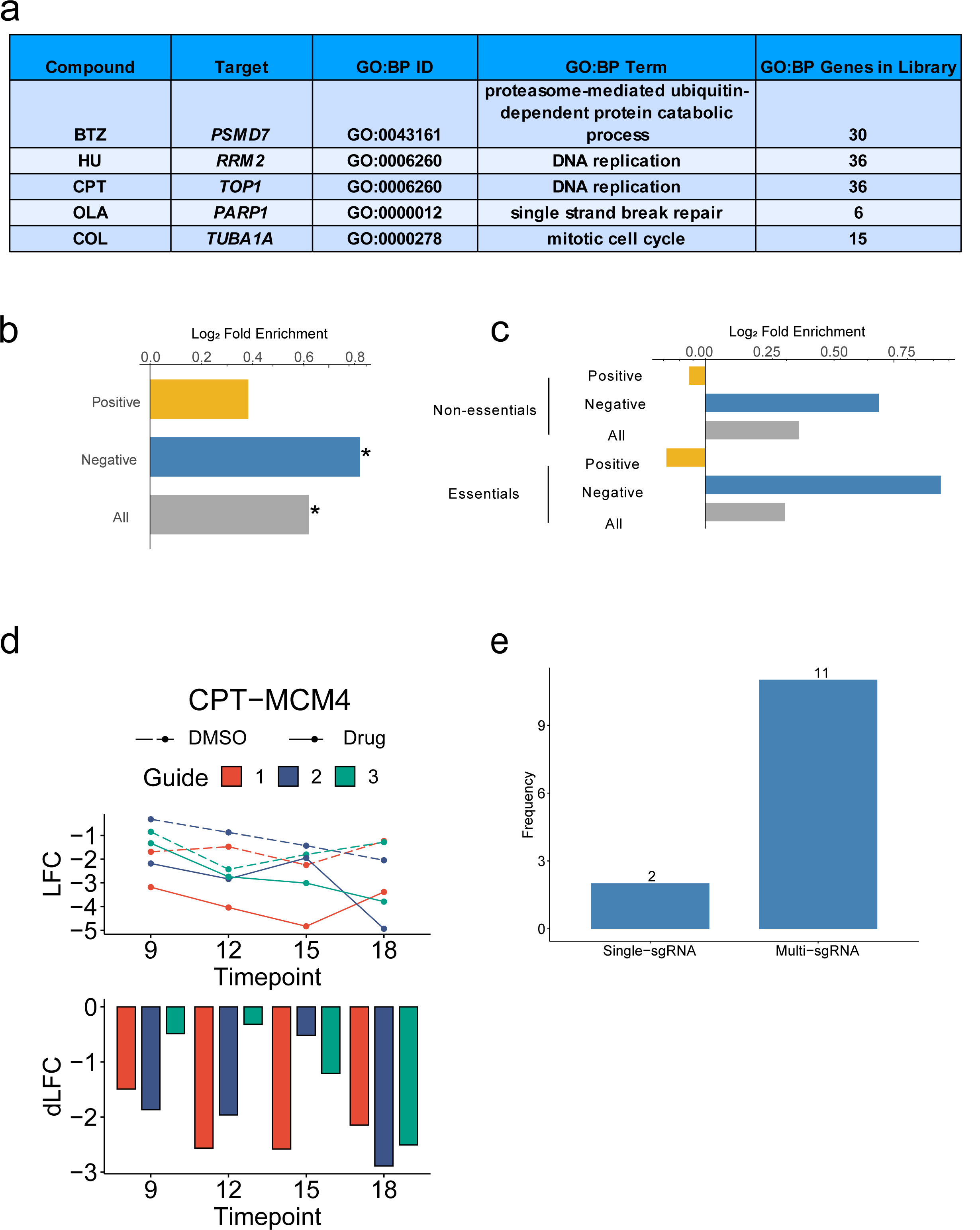
CGIs are enriched for compound mode-of-action. **a** Table of screened compounds, gene target, Gene Ontology Biological Process (GO:BP) ID, GO:BP term that best represents compound mode-of-action, and number of term-related genes found in the targeted library. **b** Enrichment of mode-of-action related genes in compound CGI pairs. X-axis: log_2_ fold enrichment. All: all CGIs, Negative: negative CGIs only, Positive: positive CGIs only. * represents p-value < 0.05. **c** Enrichment of mode-of-action related genes in compound CGI pairs for essential and non-essential genes. **d** *Top*: Guide-level LFC line plots across time points for CPT-MCM4. Orange: sgRNA 1, blue: sgRNA 2, green: sgRNA 3. Dotted line: DMSO; solid line: compound. *Bottom*: Barplot of raw differential LFC (dLFC) score for each guide. **e** Barplot categorizing 13 essential gene - compound interactions. See *Methods* for categorization approach.

The observation of CGIs for essential genes, and in particular, CGIs with essential genes related to the MoA was unexpected. In general, guides targeting essential genes drop out across the length of the screen, which is confirmed by our ROC analysis reflecting discrimination of essential genes from non-essential genes even in control conditions (Fig. 1f). We hypothesized that such interactions could be driven by rare sgRNAs that induce partial loss-of-function mutations. This hypothesis would be supported if we see only one of the three sgRNAs driving this interaction, as it is unlikely for all three sgRNAs to cause partial loss-of-function mutations in the protein encoded by the gene in question. To test this, we analyzed a total of 13 high-confidence compound - essential gene interactions found in the MoA across all compounds (Fig. S5). By quantifying whether a single sgRNA has an outlier differential log fold change (dLFC) across these 13 interactions, we determined that 11 compound-essential gene interactions were supported by multiple guides while only 2 compound-essential gene interactions were supported by a single sgRNA (Fig 3d-e). These data argue against the hypothesis of partial loss-of-function mutations induced by rare guides. An alternative explanation is that intermediate depletion of essential genes, which would occur early in the screen before wild-type protein pools are completely depleted, may result in differential phenotypes between the compound and control condition. This could result in CGIs in essential genes related to the MoA, and one would expect multiple sgRNAs targeting the same essential gene to exhibit similar phenotypes. More experiments are needed to further explore this alternative hypothesis.

### Evaluation of sensitivity and signal-to-noise characteristics of the scalable screen platform

We evaluated several aspects of the CGI hits resulting from the scalable screening platform. As a basis for our evaluations, we collected the corresponding genome-wide screen data for these compounds by either: 1) performing genome-wide screens (CPT, OLA, COL), or 2) collecting data from the genotoxin chemical screens from Olivieri et al. (HU, CPT, OLA)^12^. Raw data collected from both sources of genome-wide screens were scored for CGI hits using an adapted version of a previously described computational pipeline^23^ (see *Methods*). Table 1 shows a side-by-side comparison of the parameters for each screen source. Notably, the approach to determining compound dosage differed for each screen source, with the Olivieri screens using a lower dosage (IC_20_ determined over 12 days vs. 3 days for the scalable screens), while our genome-wide screens used a higher dosage at IC_50_. While the Olivieri screens were performed in the same cell line (hTERT-immortalized RPE-1 *TP53* knockout), our genome-wide screens were performed in HAP1 cells, a near-haploid cell line derived from the KBM7 chronic myelogenous leukemia (CML) cell line. While genetic background differences should be considered when interpreting CGIs, we reasoned that a substantial portion of CGIs should be conserved across various cell types.

#### Overlap of hits between scalable and genome-wide screens

First, we investigated whether the hits from a scalable compound screen overlapped the hits derived from its respective genome-wide screen. Hits for genome-wide screens were scored using the same computational pipeline (see *Methods*) and were defined with the same cutoffs (|CGI score| > 0.7, FDR < 0.1). Hits must point in the same direction (positive or negative in both the scalable screen and genome-wide screen) to be considered overlapping. We observed statistically significant overlap for all compounds screened (Supplementary Table 2), suggesting that the two approaches produce significantly overlapping CGI profiles.

#### Targeted library produces more hits than random subsets of genome-wide library

Next, we assessed the degree to which the 1,011 genes selected for the targeted library produced more hits than would be expected of other subsets of the genome. Specifically, we compared the number of significant CGI hits observed from the actual targeted library to 1,000 randomly selected gene sets for which we measured the number of hits observed for those genes in the corresponding genome-wide screens (as a proxy for what would have been observed had the library been constructed using each evaluated set of target genes). Each subsetted genome-wide library was rescored using a multiple hypothesis correction reflecting the reduced size (1,011 genes) to enable comparisons with our targeted library hits. As expected given our library design, the observed number of hits recovered from scalable screens generally exceeded the number of hits recovered by these randomly selected simulated libraries (Fig. S6, p < 0.004 for all compounds), suggesting the gene selection strategy (Fig. 1a) indeed biased our library towards genes with increased CGI frequency as intended for these compounds.

#### Sensitivity of scalable vs. genome-wide approach

We then compared the total yield in terms of the number of hits produced by the scalable screens as compared to their corresponding genome-wide screens (Fig. S7). On average, our genome-wide screens produced 240 significant CGIs relative to the 143 significant CGIs discovered on average across the scalable screens at the same effect size threshold and false discovery rate. For the DDR-related compounds, the number of hits for the scalable screen either exceeded the number of hits from our respective genome-wide screen (HU, OLA), or was comparable (CPT). However, this trend was reversed for colchicine, which is expected given that this DDR-focused library is likely to miss hits from a non-genotoxin. This pattern was further reflected when restricting the genome-wide screen data to the genes included in the targeted library, as the scalable screens were more sensitive to identifying hits compared to their respective genome-wide screen (except for COL). In addition, the Olivieri genome-wide screens, which were performed with much lower compound dosage, produced an average of 28.3 hits per screen (fewer than our scalable screens). This data suggests that a higher compound dosage for the screen results in a greater number of total hits.

To directly compare the sensitivity of the two sets of screens on exactly the same genes, we restricted the genome-wide screen data to only the 1,011 genes included in the targeted library (Fig. 4a). We found that the sensitivity of the scalable screens (measured as the number of hits detected relative to the total library size) was higher than the genome-wide screens on average. For example, the average hit rate for scalable screens was 14.1% compared to the hit rate of 6.6% for the genome-wide screens (after restricting to the common library genes). Furthermore, the hits unique to the scalable screens were enriched for GO terms related to the MoA for 3 of the 4 compounds compared, including all genotoxins (Supplementary Table 4). This suggested that the expanded representation per sgRNA afforded by the smaller screening format, along with sampling multiple time points across the course of the screen, enabled more sensitive detection of CGIs for this set of genes. In all but one sample (COL, T18), the hit rate for our scalable screens increased with later time sampling.

**Fig. 4.**
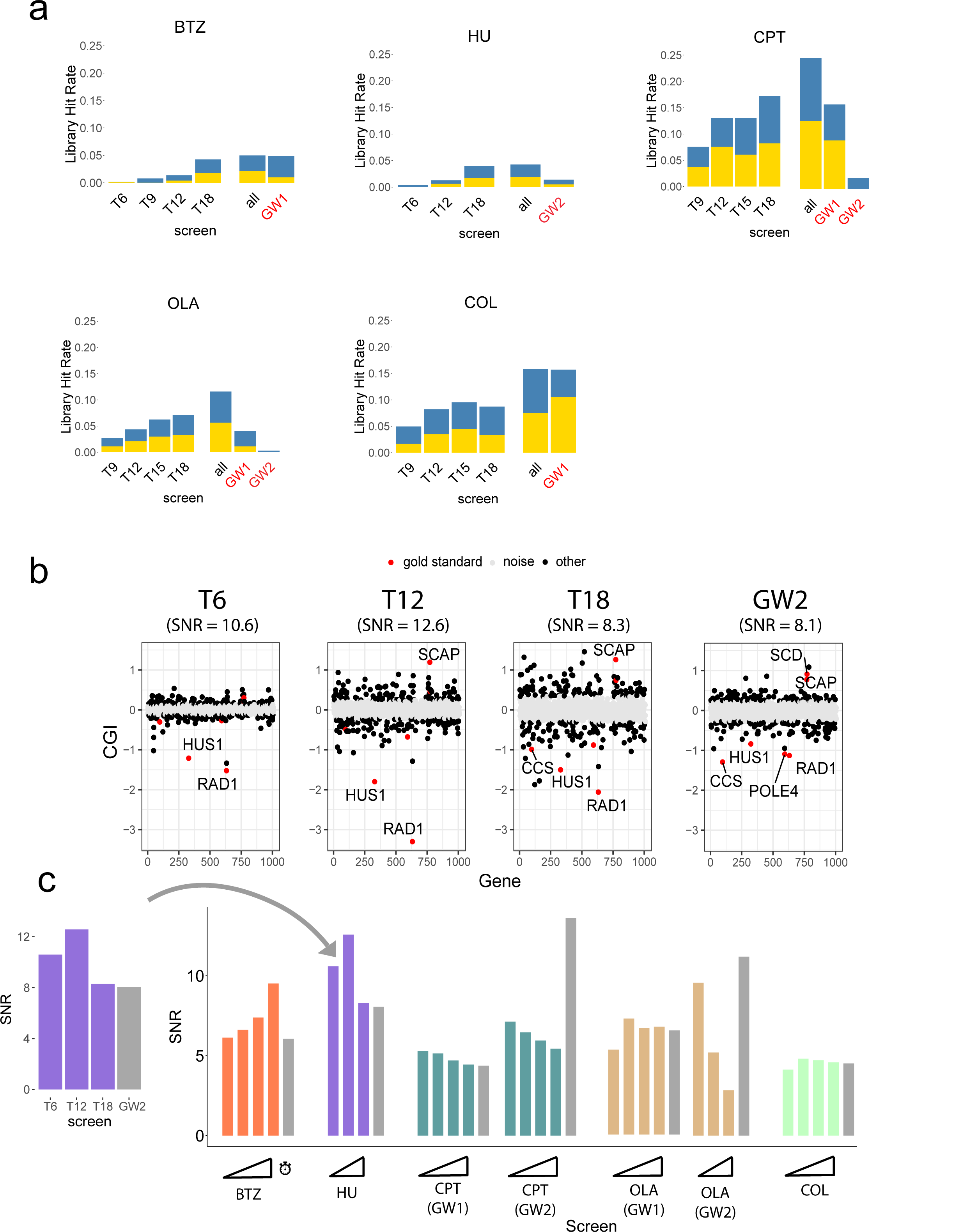
Scalable chemical screens show comparable signal-to-noise ratio. **a** Barplots of library hit rate per screen. Blue represents negative CGI hits, yellow represents positive CGI hits. For each compound, a genome-wide screen was selected for comparison (see Table 1). Red label: genome-wide screen. GW1: genome-wide screen performed for this study. GW2: genome-wide screen from Olivieri et al. All: union of hits across all time points for a given screen. **b** Signal-to-noise ratio (SNR) dotplots for HU at T6, T12, T18, as well as corresponding genome-wide screen (HU2). Genes are arranged in alphabetical order from left to right, plotted against CGI score (y-axis). Points are divided into 3 categories: 1) gold standard hits (red dots), 2) background noise (gray dots), and 3) all other genes (black dots). **c** Barplot of SNR values for all screens. SNR is defined as the mean of CGI scores (signal) divided by standard deviation of the background noise. Gray: Genome-wide screens.

#### Scalable screens have comparable signal-to-noise ratio relative to genome-wide screens

Our sensitivity analysis did not account for the identity of each CGI hit, only the total number of hits. To compare the ability of scalable vs. genome-wide screens to distinguish true hits from background noise, we developed an approach to quantify the signal-to-noise ratio (SNR). The signal was defined as the average CGI effect size across high-confidence “gold-standard” gene hits, which were formed from the intersection of each scalable screen and the corresponding genome-wide screen. The rationale in defining this gold-standard set is that hits in common between the two screening platforms are highly likely to be true positive hits and that both screen types contribute equally to forming this gold standard set such that the resulting SNR measure could be directly compared across platforms. The background noise was defined as the variance across genes with non-significant CGI effects in each assay (see *Methods* for more details). Figure 4b shows an example SNR comparison for hydroxyurea, where the SNR peaked at an intermediate time point (T12) and showed comparable SNR across all time points for the scalable screen relative to the genome-wide screen. The SNR peaked at intermediate time points during the scalable screens for multiple compounds, including HU, CPT, OLA, and COL (Fig. 4c), suggesting that the SNR is strongest at time points earlier than the typical T18 endpoint used for many published CRISPR screens. All scalable screens showed modest improvement of SNR relative to our genome-wide screens in 3 or more time points (Fig. S8). Both the CPT and OLA scalable screens showed comparable SNR to our corresponding genome-wide screen, while showing weaker SNR compared to the corresponding Olivieri screen (partially explained by the low dosage Olivieri screen, which was more sensitive to negative rather than positive CGIs). These observations generally suggest that scalable screens have comparable SNR relative to genome-wide screens. Furthermore, this SNR analysis suggests that higher SNR can frequently be achieved by sampling earlier time points than is typical for CRISPR screens in human cells (∼12 days or less) and that lower compound doses may produce chemical-genetic profiles with fewer hits but higher SNR.

### Intermediate time point CGIs reveal time-resolved dependency of multiple DNA repair pathways

To identify if certain pathways or biological processes were enriched in the CGIs of each compound screen, we performed Gene Ontology: Biological Process (GO:BP) enrichment analyses for both scalable and genome-wide screens (Supplementary Table 3, see *Methods*). For all genotoxin scalable screens (HU, CPT, OLA), we found significant enrichment (FDR < 0.2) amongst the hits in GO terms related to the compounds’ known MoA (Supplementary Table 3). The targeted library resulted in enrichment in more or a similar number of unique MoA-related GO terms relative to the genome-wide screens. In contrast, the COL scalable screen did not result in enrichment in the GO terms related to the MoA (tubulin inhibitor) whereas the genome-wide screen did. This is unsurprising given that this DDR-focused library will miss many genes in the MoA of non-genotoxins. Repeating this analysis for hits unique to the scalable screen revealed that, for genotoxins, the MoA was significantly enriched (FDR < 0.2) among these hits (Supplementary Table 4), as described above. This suggests that the scalable screens for genotoxins capture additional functionally relevant hits that were not found in the genome-wide screens.

To perform a pathway enrichment analysis more suitable for the targeted library, we manually curated 11 DNA repair and replication pathways and derived an enrichment score (see *Methods*). Genotoxins (HU, CPT, OLA) showed strong enrichment on negative CGIs for DDR and replication stress response genes, as expected (Fig. 5a). The HR pathway was strongly negatively enriched for camptothecin and olaparib, providing evidence that both compounds induced DNA breaks that employed the HR pathway for DNA repair. Interestingly, negative CGIs for camptothecin were enriched for the interstrand crosslink (ICL) repair pathway at T18 only. Colchicine negative CGIs were enriched for chromatin remodeling genes, while the positive CGIs were enriched for DNA replication, HR, and ICL pathways.

**Fig. 5.**
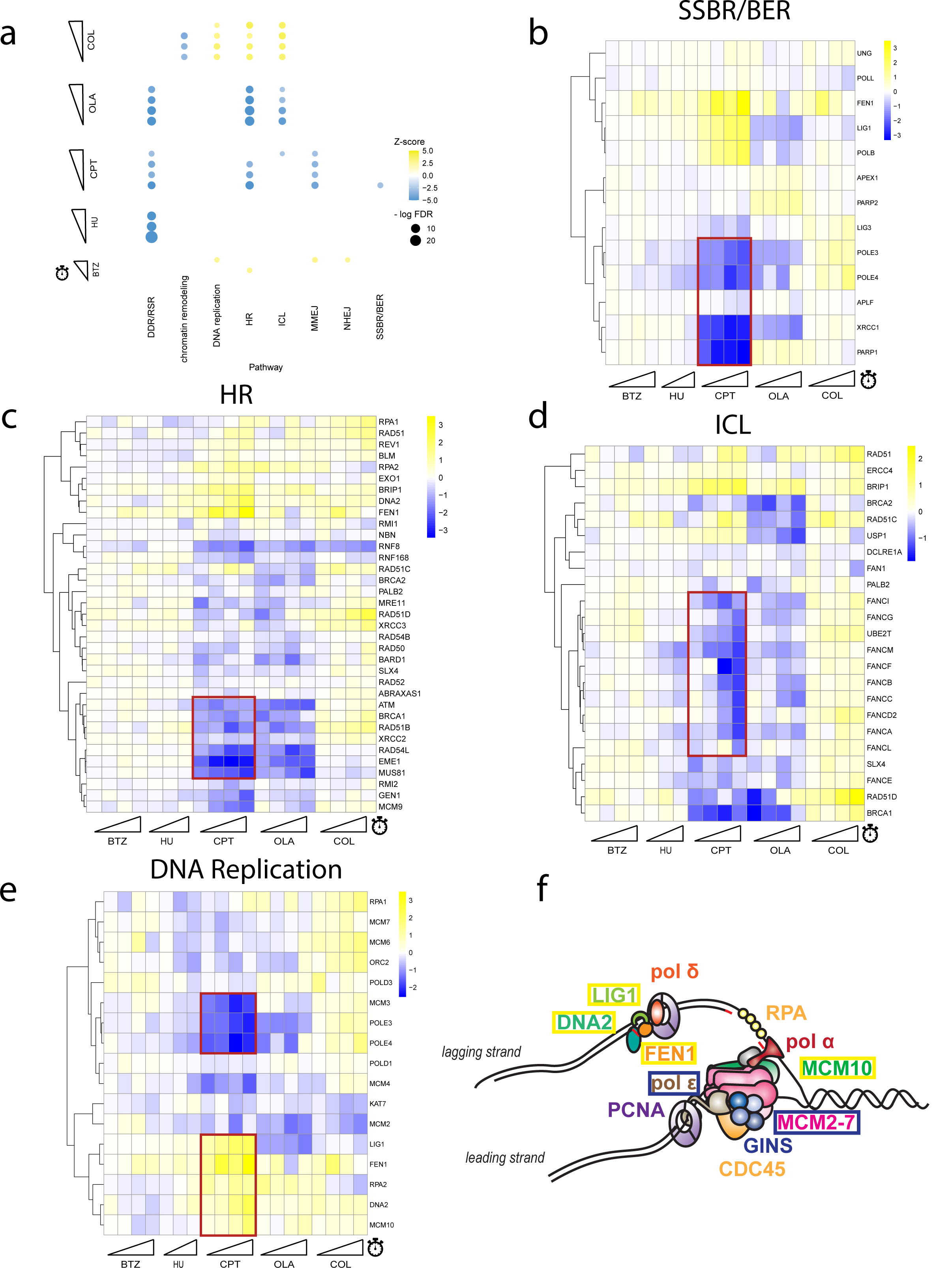
Intermediate time point data reveals time-resolution of DNA repair pathway activation. **a** Dotplot of pathway enrichment by screen. Y-axis displays each screen time point (sliding ramps represent increasing time points). X-axis displays manually curated pathways for enrichment analysis. Color indicates z-score for pathway enrichment; size of dot indicates significance (-log_10_ FDR value). Blue represents negative enrichment, yellow represents positive enrichment. Dots only appear if enrichment meets the FDR cutoff < 0.1 threshold. DDR/RSR: DNA damage response, replication stress response; HR: homologous recombination; ICL: interstrand-cross linking; MMEJ: microhomology-mediated end joining; NHEJ: non-homologous end joining; SSBR/BER: single-strand break repair, base excision repair **b** Heatmap of CGI scores for single-strand break repair (SSBR) pathway genes, using average-linkage hierarchical clustering. Blue represents negative CGI score, yellow represents positive CGI score, white represents zero CGI score. X-axis ordered by time point for each screen (sliding ramps). Relevant clusters are highlighted by the red box. **c** CGI heatmap for homologous recombination (HR) pathway. **d** CGI heatmap for interstrand-cross linking (ICL) repair pathway. **e** CGI heatmap for DNA replication genes. **f** Model showing the lagging and leading strand replication proteins. Genes outlined in yellow boxes displayed positive CGIs with CPT in **e**; genes outlined in blue boxes displayed negative CGIs with CPT in **e**.

Heatmaps of the CGI scores across the compound screens for each specific curated pathway reveal time resolution dependency on DNA repair pathways in response to camptothecin-induced damage (Fig. 5b-d, S9). DNA damage recognition proteins, such as XRCC1 and PARP1, sense and bind to sites of DNA damage before DNA repair begins with DNA polymerase activity. This is supported by the consistent strong negative CGIs of *XRCC1* and *PARP1* across all time points for the camptothecin screen, and strong negative CGIs with *POLE3/4* only at later time points (Fig. 5b). Inspection of the HR genes (*ATM, BRCA1, RAD51B, XRCC2, RAD54L, EME1, MUS81*) revealed expected negative CGIs with camptothecin (Fig. 5c). Interestingly, many members of the ICL pathway (*FANCI, FANCG, FANCM, FANCF, FANCB, FANCC, FANCD2, FANCA*) showed strong negative interactions with camptothecin at later time points (T15, T18) only (Fig. 5d). This suggests that members of the ICL pathway may recognize an ICL-like intermediate complex and serve as an alternative repair pathway mechanism for double strand breaks induced by camptothecin that activates after the initial HR pathway response. The intermediate time point CGI data has the ability to capture time-resolution data on DNA repair pathways, potentially revealing the sequence in which different DNA repair pathways respond to DNA damage.

Lagging and leading strand genes showed opposing interaction patterns due to delayed replication caused by camptothecin trapping topoisomerase I on DNA strands (Fig. 5e-f). *POLE3* and *POLE4*, which displayed strong negative interactions at late time points for camptothecin, encode subunits of DNA polymerase epsilon, which synthesizes the leading strand during replication. In contrast, *FEN1*, *DNA2*, *LIG1*, *MCM10*, which all displayed strong positive interactions with camptothecin, encode proteins that act on the lagging strand. The deletion of these genes is thought to delay Okazaki fragment processing, slowing DNA synthesis^34^. Given that camptothecin creates DNA:topoisomerase adducts, slowed DNA synthesis decreases the probability that the replisome machinery will encounter these adducts, which may explain the improved cell viability compared to non-treated cells. Clustering of CGI scores in these curated pathways can provide evidence for distinct biological roles of protein complexes or specific biological pathways.

## Discussion

Motivated by previous efforts that established scalable CGI profiling platforms in *S. cerevisiae*^5^, we developed and characterized a small, DDR-focused library for CRISPR screens. Our library consists of 3,033 experimentally validated guides targeting 1,011 genes, and thus is approximately 1/20th the size of typical genome-wide libraries. Screening with this library requires substantially fewer reagents, both in terms of cell culture and sequencing, to maintain a sufficient representation and provide quantitative measures of CGIs. The reduction of tissue culture plates afforded by this scalable approach enables higher coverage (1000X representation of each sgRNA), greater time point resolution of CGIs, and larger number of technical replicates for added statistical power when determining significant CGIs. Based on 5 proof-of-principle screens, we found that it provides increased sensitivity to interactions for the compressed gene space at a comparable or better SNR than genome-wide screens.

One important practical advantage to a scalable screening platform like the one we presented here is the cost-efficiency of sampling interactions at multiple time points. The temporal resolution of CGI data has not been previously explored and may provide novel insights into how biological pathways respond to chemical perturbations over time. We found that our platform could detect the sequential action of DNA damage recognition (*XRCC1/PARP1*) before DNA repair (*POLE3/4*) from the CGI profile of the camptothecin screen. In addition, we found strong negative CGIs between camptothecin and Fanconi anemia complementation group (FANC) genes at later time points only, suggesting a delayed dependency on the ICL repair pathway in response to DNA damage induced by camptothecin. To our knowledge, this potential switch from HR/SSBR to ICL response to camptothecin has not been previously described.

There are notable limitations to our approach. First, given the limited gene space covered by the targeted library (1,011 genes), there are many CGIs that could be informative about compounds’ MoA that will be missed. Indeed, we found that simply performing functional enrichment analysis on the resulting hits can be substantially less informative for a small library as compared to a genome-wide screen in which the entire genome is targeted (Supplementary Table 3) for some compounds (e.g. Colchicine). In our case, this library is enriched for genes involved in DDR, so the platform is highly resolved for compounds with DDR-related MoA, but will be less powerful for compounds targeting other functions. Future work could focus on developing similar targeted libraries designed to capture other bioprocesses. A second limitation of the screening platform we describe here is that, since the library design was completed, a wealth of additional data from CRISPR screens has become publicly available (e.g. the DepMap project has substantially expanded^35^). Future library design efforts should leverage all the latest available screening data, which we expect would improve the extent to which the resulting profiles are representative of genome-wide profiles.

In general, chemical-genetic screens provide a powerful lens for characterizing novel compounds and identifying new therapeutic opportunities for drugs already in use. However, the space on which CGI technology could be productively applied is enormous. There are hundreds of large compound libraries, including both naturally occurring and synthetic compounds, in addition to the large space of clinically approved drugs. Furthermore, exploring the functional impacts of combinatorial drug treatments is also of interest. In addition to the large chemical space, the cell type context in which CGI screens are conducted is also important. We focus on RPE-1 and HAP1 cells here, but screening a variety of cell types, especially those well-matched to specific biological or therapeutic questions, will be important. Scalable screening platforms that enable rapid application of chemical-genetic screens across all these critical dimensions will play an important role in realizing the full potential of this technology for drug discovery.

## Methods

### Cell lines and culture conditions

RPE-1 hTERT Cas9 *TP53^-/-^* (female human hTERT-immortalized retinal pigmented epithelial cells) was constructed as previously described^22^. RPE-1 cells were grown in Dulbecco’s Modified Eagle Medium/Nutrient Mixture F-12 (DMEM:F12) supplemented with 10% FBS and 1% penicillin-streptomycin. Cells were grown at 37°C and 5% CO_2_ in standard tissue culture incubators. Cells were regularly tested for mycoplasma contamination with the PCR-based Venora GeM Mycoplasma Detection Kit; no mycoplasma contamination was detected during this study.

### Targeted CRISPR library design

Selection of genes for the compressed, targeted CRISPR library aimed to target the ∼1,000 protein encoding genes that were likely to display variable cell fitness effects. Four categories were chosen to select those genes: 1) 349 DNA damage response genes, 2) 100 genes capturing the most variance across published CRISPR screens, 3) 216 genes that captured subtle fitness defects across time-course CRISPR screen data, and 4) 463 genes that have a high degree of genetic interactions. Category 2 to 4 genes were selected from genome-wide data sets, but the 684 core essential genes defined in Hart et al 2017^18^ were excluded from each selection process.

Category 1 genes were manually curated and selected by DDR field experts. In contrast to categories 2 to 4, genes were included regardless of their essentiality status. Category 2 genes were selected by extracting CRISPR screen data from the major genome-wide cell fitness readout data sets available at the time of library generation. Overall, those comprised 61 cell lines^36–40^. Raw read count data were downloaded from the GenomeCRISPR database^41^. Gene essentiality scores (Bayes Factors) for each screen were computed using the Bagel pipeline^42^, followed by batch correction using the combat method implemented in the sva Bioconductor package in R^43^ (*Surrogate Variable Analysis*. R package version 3.48.0). The top 100 genes with the greatest average variance across batch-corrected fitness scores were selected to constitute category 2 genes. Category 3 genes were selected by using time-course genome-wide CRISPR screen data from HAP1 cells. To obtain robust temporal sgRNA dropout patterns, the data of seven HAP1 TKOv3 library screens, of which three had intermediate time points that were taken every three days up to the endpoint measurement (T18)^44^, were merged. The consensus log_2_ fold-change (T[3 – 18] / T0) was computed for each sgRNA at each time point. To classify genes by their dropout pattern, we defined distinct short time-series expression miner (STEM) clusters for all ∼71k sgRNAs that captured subtle fitness defect changes over the length of the screen. Overall, we defined 12 distinct clusters. To assign a gene to a cluster, we then only kept genes where two (of the four) independent sgRNAs were clustered and no other cluster contained more than a single sgRNA targeting that gene. We then selected even numbers of genes from each cluster for the compressed library. Category 4 genes were selected from an unpublished genome-wide genetic interaction data set measured in HAP1 cells. Specifically, genome-wide CRISPR-Cas9 screens had been performed with the TKOv3 library in HAP1 wildtype (control) and HAP1 knockout cells in which a specific knockout was introduced. Overall, 33 genetic backgrounds were screened at the time of the library design. Quantitative GI (qGI) scores were extracted from those 33 screens^45^, and the 463 most frequent interacting genes at a qGI-associated FDR of 10% were chosen.

Overall, 1,011 total genes were selected for the targeted library, with several genes overlapping multiple categories (see Supplementary Table 5 for complete list). For 990 genes, we selected the 3 best performing guides from the genome-wide TKOv3 library. Those were defined based on a comprehensive set of screens performed in 33 distinct genetic backgrounds in HAP1 cells. Specifically, we quantified genetic interactions between each gene in the TKOv3 library with the defined background mutation present in a given HAP1 clone. To measure sgRNA quality, we utilized the sgRNA genetic interaction scores by computing the pairwise Pearson correlation coefficients (PCC) between all sgRNA targeting the same gene across their genetic interaction profiles. Per sgRNA, the PCCs were summed up and the sgRNA with the three highest scores were chosen. The remaining 21 genes were not found in the TKOv3 library and were manually chosen for the targeted library. In total, there are 3,033 sgRNAs targeting 1,011 genes in the targeted CRISPR library.

This custom, DDR-focused targeted library was constructed by Cellecta, with each sgRNA cloned into the pRSG16-U6-sg-HTS6C-UBiC-TagRFP-2A-Puro plasmid. The plasmid contains a puromycin-resistance cassette for selection of cells that contain a library sgRNA during the pooled screen.

### Proof-of-principle scalable CRISPR-Cas9 chemical screens

A detailed protocol of the scalable CRISPR chemical screens can be found here^22^. The major steps are briefly described below.

#### Compound concentration determination

Compounds were all diluted in vehicle (DMSO) in preparation for screening purposes. To determine the compound dosage used for each screen, we conducted an ATPase cell viability assay. RPE-1 hTERT cells were initially seeded on day 1 with a density of 1,500 cells per well in 96-well plates. On day 2, media was removed and replaced with either media + compound in a range of 10 or more doses, or with vehicle control (0.5 % w/v DMSO), in triplicates. Cells were incubated for 72 hours, and on day 5, CellTiterGloⓇ luminescent assay (Promega #G75752) was used to approximate cell viability and generate a dose-response curve. Luminescence intensities were measured on a Promega GloMax Microplate Reader. The relative survival of compound-treated vs. untreated cells was expressed as a percentage of the untreated DMSO control. For each compound, a dose corresponding to IC_20_ (20% growth inhibition relative to DMSO controls) was selected for screening. Before initiating a screen, the dosage effect was verified in the 15-cm tissue culture plates that would be used for the screen.

#### Lentivirus production and infection

Cells in 15-cm dishes at 70% confluency were transfected with 1.9×10^9^ TU/mL of lenti-library and 10 ug/mL polybrene, yielding a MOI of 0.2 (1 in 5 cells infected). A separate 15-cm control plate of cells was cultured in parallel. 24 hr after transfection, the medium was replaced with fresh medium containing 3 ug/mL puromycin to transduced plates and to the control plate. 48h after puromycin treatment, cells completely died in the control plate. The remaining cells in the transduced plates, which have all presumably integrated a sgRNA, are pooled and pelleted.

#### Pooled screen

At T0, cells were split into media with vehicle control (DMSO) or with one of the 5 compounds at an IC_20_ dosage, seeding ∼ 3×10^6^ cells per replicate (1 15 cm plate per replicate) at a desired 1,000-fold sgRNA coverage. Additionally, cell pellets were collected at T0. Cells were split every 3 days into a combination of new medium and compound or DMSO, ensuring 1,000-fold sgRNA coverage at each split. Cell pellets were also collected every 3 days until T18 (T3, T6, T9, T12, T15, T18). Technical replicates were independent throughout the screen (cells were not pooled together after each passage).

#### NGS library prep

Genomic DNA was extracted from each cell pellet using the Promega Wizard Genomic DNA Purification Kit (Promega #A1120), following standard protocol. Next, two-round PCR was performed using the Cellecta NGS prep kit for sgRNA barcode libraries in pRSG16/17 (KOHGW) (Cellecta # LNGS-120) and the Supplementary Primer Sets (Cellecta #LNGS-120-SP) to amplify the sgRNA and append Illumina sequencing adapters and index barcodes for each replicate sample. We used 20 ug of genomic DNA in 50 uL 1st-round PCR reaction volume, and 5 uL of PCR1 product for 50 uL 2nd-round PCR reaction volume. QIAquick PCR Purification (Qiagen #28104) and Gel Extraction Kits (Qiagen #28704) were used to clean up the library prep, and samples were run on a 2% agarose-1X TAE gel to check product size before next-generation sequencing. A maximum of 48 samples were pooled and sequenced on a single lane on the Illumina NextSeq 550 (standard Single-Read 150-cycles) at the UMGC (University of Minnesota Genomics Center) using common sequencing primers provided by UMGC and indexing primers provided by Cellecta.

### Genome-wide screens

Genome-wide screens were conducted in a similar fashion to screens described here^45^. These screens utilized the Toronto KnockOut version 3 (TKOv3) genome-wide library^18^ in the near-haploid HAP1 cell line. Each compound was screened at an IC_50_ concentration, and library representation was maintained at ∼250-fold coverage. Cell pellets were collected and sequenced at T0 and T18 for all compounds (except T13 for olaparib).

CRISPR genome-wide screen data was not available for bortezomib. Instead, CGI hits were derived from a bortezomib shRNA screen^46^. The shRNA library used for this screen targeted 7,712 genes involved in proteostasis, cancer, apoptosis, kinases, phosphatases, and drug targets^46^. This screen data was used for comparison vs. the scalable bortezomib screen.

### Raw read counts

Demultiplexed FASTQ files were generated using the Illumina bcl2fastq software. These files were used as input for the Cellecta “NGS Demultiplexing and Alignment Software,” along with a “Sample Description File” that matched index barcode to each sample and a “Library Configuration File” containing a list of target sgRNA guide sequences. The Cellecta software generated a table of raw read counts for each sgRNA (row) and each sample (column).

### Chemical-genetic interaction scoring

CGIs were scored using an adapted version of the Orthrus software^23^. Raw read counts were normalized by read depth for each sample. Per-guide-level log_2_ fold changes (LFC) were calculated between an intermediate or end time point and starting time point (T0). LFC values underwent two additional normalization steps: 1) MA-transformation, where guide-level ratios (M) were plotted against mean average (A) guide-level LFC data, and 2) loess (**<u>l</u>**ocally **<u>es</u>**timated **<u>s</u>**catterplot smoothing) regression, which bins the data with equal bin sizes along the A values and fits a smooth curve through the data points within each bin. Replicate normalized LFC values are averaged before downstream steps. Then, the guide-level CGI scores were derived from calculating the differential normalized LFC values between compound and control screens. Guide-level CGI scores per gene were averaged and tested for significance using the moderated t-test from the *limma* R package^24^. P-values were adjusted through Benjamini-Hochberg multiple testing correction per screen. The code for CGI scoring is available at this link.

The CGI scoring approach described above was used to derive CGI hits from raw read count screen data for the scalable screens, for the genome-wide screens performed by us, and for the genome-wide screens performed by Olivieri et al.

### Screen quality control metrics

To assess the quality of the resulting CRISPR screen data, we used three quality control metrics: 1) replicate correlation on LFC and CGI scores, 2) core essential gene dropout, and 3) within-vs-between context correlation (WBC) scores. Replicate correlation was computed with a Pearson’s correlation coefficient on the vector of LFC values between all possible replicate pairs (AB, AC, BC). Using a core essential gene standard defined by the Broad Dependency Map (DepMap^35^) data (genes observed to be broadly essential across many cell lines, see *Essential genes analysis*), we generated AUC-ROC (area under the curve – receiver operating characteristic) values to quantify how well core essential genes drop out relative to non-essential genes throughout the length of the screen. WBC scores were calculated as previously described^25^.

### Visualization of clustering analyses

The heatmap for Fig. 2a was generated using a Pearson correlation coefficient similarity metric and average-linkage, hierarchical clustering and visualized in Java TreeView version 1.2.0. The heatmaps for Fig. 4 were generated using Pearson correlation coefficient similarity and average-linkage, hierarchical clustering options from the *pheatmap* R package.

### Library hit rate

Library hit rate is defined as the ratio of the number of significant hits to the library size (in number of genes). To compare library hit rates from scalable vs. genome-wide screens, we subset the 1,011 targeted library genes from the genome-wide screen library and applied multiple hypothesis correction only on this subset to avoid penalizing the genome-wide screen data for additional tests beyond the 1,011 genes. Next, we tallied the number of total CGIs, positive CGIs, and negative CGIs for each screen, using a CGI score cutoff of greater than 0.7 (for positive CGIs) or less than −0.7 (for negative CGIs) and a false discovery rate (FDR) lower than 10% (Supplementary Table 1). We then calculated the library hit rate for each time point scalable screen vs. genome-wide screen (Fig. 3a). We also took the union of unique gene hits across all time points for the scalable screens to simplify comparisons to genome-wide screens.

### Signal-to-noise ratio

For each screen, each of the 1,011 genes in the targeted sgRNA library is associated with a CGI score and FDR value. A “significant hit” is defined by a CGI cutoff (|CGI| > 0.7) and FDR cutoff (FDR < 0.1). Genes are divided into 3 categories: 1) “gold standard” hits, defined by intersecting the hits from a scalable time point screen with genome-wide screen hits, 2) the “background noise” set, defined as genes with CGI values in the middle 80% of the distribution of CGI values (10th-90th percentile, expected to reflect random variation across non-interacting genes), and 3) all other genes. We defined the signal-to-noise ratio as follows:

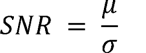

where *μ* is the average CGI score across all gold standard genes, and *σ* is the standard deviation of the CGI scores of genes in the background noise set. SNRs were calculated individually for each time point of a given compound screen.

### Mode-of-action fold enrichment analysis

To quantify to what extent the known mode-of-action (MoA) of a compound is enriched in its CGI profile, we perform the following analysis. First, we select a Gene Ontology: Biological Process (GO:BP) term that best describes the biological process or pathway perturbed by the compound in question. This GO:BP term must have the gene encoding the protein target of the compound annotated to it. Second, we define the fold enrichment (FE) metric for each time point screen with the following equation:

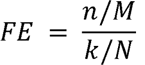

where n is the number of hits found in the GO:BP term ascribed to the MoA, M is the number of significant hits for that time point of the screen, k is the number of library genes found to be annotated to the GO:BP MoA term, and N is the total number of library genes. The same equation can be used to describe the global fold enrichment across all screens, where n is the number of compound-hit pairs found in the MoA GO:BP for each compound, M is the number of compound-hit pairs detected, k is the number of compound-library gene pairs found to be annotated to the GO:BP MoA term, and N is the total number of compound-library gene pairs. The global fold enrichment metric can further be broken down by considering essential and non-essential genes separately (see “Essential gene analysis” section), or negative or positive CGIs only. Each FE metric is reported in log_2_ transformation and associated with a p-value calculated from a hypergeometric test.

### Essential gene analysis

An essential gene standard is defined from the CRISPR screen Broad Dependency Map (DepMap^35^) 20Q2 dataset. A gene is defined as essential if it exhibits a < -1 CERES score in > 60% of the 769 DepMap cancer cell lines^35^. The targeted library contains 55 essential genes, and this essential gene set was used to generate AUC-ROC curves to assess screen quality.

For the essential gene mode-of-action analysis, we derived the following approach to determine if a CGI was supported by single or multiple sgRNAs: 1) for each gene, calculate residuals to average DMSO LFC at each time point for each of the 3 sgRNAs, 2) calculate the standard deviation of these residuals, 3) calculate an uncorrected differential score matrix between compound and DMSO LFC, 4) determine if each differential score exceeds one standard deviation threshold in at least two or more time points of a screen, and 5) determine whether the differential score is supported by 1 or multiple of the 3 targeting sgRNAs per gene.

### Pathway enrichment analysis

For GO:BP enrichment analysis, we used the “enrichGO” function in the *clusterprofiler* R package. Parameters include a p-value cutoff at 0.05, a q-value cutoff at 0.2, minimum gene set size at 5, and maximum gene set size at 200. P-values were adjusted by the Benjamini-Hochberg method.

For the targeted library only enrichment analysis, a set of 11 DNA repair and replication pathways were manually curated for the targeted library genes (see Supplementary Table 6). Assuming a normal distribution for the CGI scores, a z-score was calculated using the following formula for each pathway:

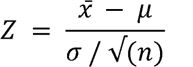

where *x* = average CGI score of all genes annotated to the pathway, *μ* = the average CGI score across all library genes, *σ* = the standard deviation of CGI scores across all library genes, and *n* = number of genes annotated to the pathway. A two-tailed p-value is calculated for each z-score using the pnorm function in R. The p-values are then adjusted for multiple comparisons using the Benjamini-Hochberg method.

## Supplementary Information

**Fig. S1 Effect of dosage on compound screen hits. a** Scatterplot of control (DMSO) fitness vs. compound fitness (log_2_ fold change) for T6 BTZ screen at IC_50_ dosage. Negative and positive chemical-genetic interactions (CGIs) are indicated in blue and yellow, respectively. Each point represents a gene. The top 5 negative/positive hits are labeled. **b** Scatterplot of DMSO fitness vs. T12 BTZ fitness (IC_50_). **c** Scatterplot of DMSO fitness vs. T6 BTZ fitness (IC_20_). **d** Scatterplot of DMSO fitness vs. T12 BTZ fitness (IC_20_).

**Fig. S2 Correlation matrix for CGI scores.** Pearson’s correlation coefficient matrix on CGI scores for each screen. Positive correlations are represented in shades of blue; negative correlations are represented in shades of red. Sliding ramp represents increasing time points.

**Fig. S3 Volcano plots for all compound screens.** Volcano plot for each compound screen (BTZ = bortezomib, HU = hydroxyurea, CPT = camptothecin, OLA = olaparib, COL = colchicine). Negative and positive CGIs are indicated in blue and yellow, respectively. Each point represents a gene. False discovery rate (FDR) values were estimated using the Benjamini-Hochberg method. Cutoffs for significant CGIs (hits) were set at FDR = 0.1 and |CGI| > 0.7 (gray dashed lines). The top five negative and positive hits are labeled.

**Fig. S4 LFC scatterplots for all compound screens.** Scatterplot of control (DMSO) fitness vs. compound fitness (BTZ = bortezomib, HU = hydroxyurea, CPT = camptothecin, OLA = olaparib, COL = colchicine). Negative and positive CGIs are indicated in blue and yellow, respectively. Each point represents a gene. The top five negative and positive hits are labeled.

**Fig. S5 Essential gene analysis plots for all essential gene - compound pairs.** *Top*: Guide-level LFC line plots across time points for each compound-essential gene pair. Orange: sgRNA 1, blue: sgRNA 2, green: sgRNA 3. Dotted line: DMSO; solid line: compound. *Bottom*: Barplot of raw differential LFC (dLFC) score for each guide across time points.

**Fig. S6 Random distribution of total significant hits recovered from simulation of compressed gene library.** 1,000 simulations of hit rate for randomly selected 1,011-gene library based on subsets of the genome-wide library. A separate distribution was generated for each compound. Red line: the observed number of significant hits from each respective scalable screen. Genome-wide screen data generated for this study is denoted with “1” following compound name in title. Genome-wide screen data generated by Olivieri et al. is denoted with “2” following compound name in title.

**Fig. S7 Barplots of number of hits per compound screen**. Red: scalable screen. Blue: Genome-wide screen. Green: Genome-wide screen restricted to genes in targeted library (1,011 genes). “1” denotes genome-wide screen performed for this study. “2” denotes genome-wide screen from Olivieri et al.

**Fig. S8 SNR plots for all compound screens.** Signal-to-noise ratio (SNR) dotplots. Genes are arranged in alphabetical order from left to right (x-axis), plotted against CGI score (y-axis). Points are divided into 3 categories: 1) gold standard (red dots), 2) background noise (gray dots), and 3) all other genes (black dots). **a** SNR dotplot for bortezomib (at IC_20_ dose). **b** SNR dotplot for bortezomib (at IC_50_ dose). **c** SNR dotplot for camptothecin (vs. our genome-wide screen). **d** SNR dotplot for camptothecin (vs. genome-wide screen from Olivieri et al) **e** SNR dotplot for olaparib (vs. genome-wide screen conducted for this study). Note that comparison vs. genome-wide screen from Olivieri et al. was not included because there were not enough hits from the genome-wide screen to conduct SNR analysis. **f** SNR dotplot for colchicine (vs. genome-wide screen conducted for this study).

**Fig. S9 CGI heatmap for all other curated pathways. a** Heatmap of CGI scores for DNA damage response / replication stress response (DDR/RSR) pathway genes, using average-linkage hierarchical clustering. Blue represents negative CGI score, yellow represents positive CGI score, white represents zero CGI score. X-axis ordered by screen time point (sliding ramps). **b** Heatmap for chromatin remodeling genes. **c** Heatmap for microhomology-mediated end joining (MMEJ) pathway. **d** Heatmap for mismatch repair (MMR) pathway. **e** Heatmap for nucleotide excision repair (NER) pathway. **f** Heatmap for non-homologous end joining (NHEJ) pathway. **h** Heatmap for translesion synthesis (TLS) genes.

**Supplementary Table 1: Significant hits list for each scalable compound screen**

**Supplementary Table 2: Number of hits and overlapping hits for each screen**

**Supplementary Table 3: GO:Biological Process enrichment for each screen**

**Supplementary Table 3: GO:Biological Process enrichment on hits unique to scalable screens**

**Supplementary Table 5: Categorization of targeted library genes and guide sequences**

**Supplementary Table 6: Manual curation of 11 DDR-related pathways**

## References

1. Colic, M. & Hart, T. Chemogenetic interactions in human cancer cells. Comput. Struct. Biotechnol. J. 17, 1318–1325 (2019).

2. Topatana, W. et al. Advances in synthetic lethality for cancer therapy: cellular mechanism and clinical translation. J. Hematol. Oncol.J Hematol Oncol 13, 118 (2020).

3. Tong, A. H. Y. et al. Systematic Genetic Analysis with Ordered Arrays of Yeast Deletion Mutants. Science 294, 2364–2368 (2001).

4. Hillenmeyer, M. E. et al. The Chemical Genomic Portrait of Yeast: Uncovering a Phenotype for All Genes. Science 320, 362–365 (2008).

5. Piotrowski, J. S. et al. Functional Annotation of Chemical Libraries across Diverse Biological Processes. Nat. Chem. Biol. 13, 982–993 (2017).

6. Mohr, S., Bakal, C. & Perrimon, N. Genomic Screening with RNAi: Results and Challenges. Annu. Rev. Biochem. 79, 37–64 (2010).

7. Jinek, M. et al. A Programmable Dual-RNA–Guided DNA Endonuclease in Adaptive Bacterial Immunity. Science 337, 816–821 (2012).

8. Ran, F. A. et al. Genome engineering using the CRISPR-Cas9 system. Nat. Protoc. 8, 2281–2308 (2013).

9. Shalem, O. et al. Genome-Scale CRISPR-Cas9 Knockout Screening in Human Cells. Science 343, 84–87 (2014).

10. Wang, T., Wei, J. J., Sabatini, D. M. & Lander, E. S. Genetic screens in human cells using the CRISPR-Cas9 system. Science 343, 80–84 (2014).

11. Ruiz, S. et al. A Genome-wide CRISPR Screen Identifies CDC25A as a Determinant of Sensitivity to ATR Inhibitors. Mol. Cell 62, 307–313 (2016).

12. Olivieri, M. et al. A Genetic Map of the Response to DNA Damage in Human Cells. Cell 182, 481–496.e21 (2020).

13. Olivieri, M. & Durocher, D. Genome-scale chemogenomic CRISPR screens in human cells using the TKOv3 library. STAR Protoc. 2, 100321 (2021).

14. Hundley, F. V. et al. A comprehensive phenotypic CRISPR-Cas9 screen of the ubiquitin pathway uncovers roles of ubiquitin ligases in mitosis. Mol. Cell 81, 1319–1336.e9 (2021).

15. Hundley, F. V. & Toczyski, D. P. Chemical-genetic CRISPR-Cas9 screens in human cells using a pathway-specific library. STAR Protoc. 2, 100685 (2021).

16. Jackson, S. P. & Bartek, J. The DNA-damage response in human biology and disease. Nature 461, 1071–1078 (2009).

17. Esmaeili Anvar, N., et al. Combined genome-scale fitness and paralog synthetic lethality screens with just 44k clones: the IN4MER CRISPR/Cas12a multiplex knockout platform. BioRxiv Prepr. Serv. Biol. 2023.01.03.522655 (2023) doi:10.1101/2023.01.03.522655.

18. Hart, T. et al. Evaluation and Design of Genome-Wide CRISPR/SpCas9 Knockout Screens. G3 Genes Genomes Genet. 7, 2719–2727 (2017).

19. Haapaniemi, E., Botla, S., Persson, J., Schmierer, B. & Taipale, J. CRISPR-Cas9 genome editing induces a p53-mediated DNA damage response. Nat. Med. 24, 927–930 (2018).

20. Brown, K. R., Mair, B., Soste, M. & Moffat, J. CRISPR screens are feasible in TP53 wild-type cells. Mol. Syst. Biol. 15, e8679 (2019).

21. Haapaniemi, E., Botla, S., Persson, J., Schmierer, B. & Taipale, J. Reply to “CRISPR screens are feasible in TP53 wild-type cells”. Mol. Syst. Biol. 15, e9059 (2019).

22. Lin, K. et al. Scalable CRISPR-Cas9 chemical genetic screens in non-transformed human cells. STAR Protoc. 3, (2022).

23. Ward, H. N. et al. Analysis of combinatorial CRISPR screens with the Orthrus scoring pipeline. Nat. Protoc. 16, 4766–4798 (2021).

24. Ritchie, M. E. et al. limma powers differential expression analyses for RNA-sequencing and microarray studies. Nucleic Acids Res. 43, e47 (2015).

25. Billmann, M. et al. Reproducibility metrics for context-specific CRISPR screens. Cell Syst. 14, 418–422.e2 (2023).

26. Lee, J. & Dunphy, W. G. Rad17 Plays a Central Role in Establishment of the Interaction between TopBP1 and the Rad9-Hus1-Rad1 Complex at Stalled Replication Forks. Mol. Biol. Cell 21, 926–935 (2010).

27. Parrilla-Castellar, E. R., Arlander, S. J. H. & Karnitz, L. Dial 9–1–1 for DNA damage: the Rad9–Hus1–Rad1 (9–1–1) clamp complex. DNA Repair 3, 1009–1014 (2004).

28. Rose, M., Burgess, J. T., O’Byrne, K., Richard, D. J. & Bolderson, E. PARP Inhibitors: Clinical Relevance, Mechanisms of Action and Tumor Resistance. Front. Cell Dev. Biol. 8, (2020).

29. Pommier, Y. Topoisomerase I inhibitors: camptothecins and beyond. Nat. Rev. Cancer 6, 789–802 (2006).

30. Bian, L., Meng, Y., Zhang, M. & Li, D. MRE11-RAD50-NBS1 complex alterations and DNA damage response: implications for cancer treatment. Mol. Cancer 18, 169 (2019).

31. Ciccia, A., Constantinou, A. & West, S. C. Identification and Characterization of the Human Mus81-Eme1 Endonuclease*. J. Biol. Chem. 278, 25172–25178 (2003).

32. Taylor, E. W. THE MECHANISM OF COLCHICINE INHIBITION OF MITOSIS : I. Kinetics of Inhibition and the Binding of H3-Colchicine. J. Cell Biol. 25, 145–160 (1965).

33. Margueron, R. & Reinberg, D. The Polycomb complex PRC2 and its mark in life. Nature 469, 343–349 (2011).

34. Zheng, L. & Shen, B. Okazaki fragment maturation: nucleases take centre stage. J. Mol. Cell Biol. 3, 23–30 (2011).

35. Tsherniak, A. et al. Defining a Cancer Dependency Map. Cell 170, 564–576.e16 (2017).

36. Wang, T. et al. Identification and characterization of essential genes in the human genome. Science 350, 1096–1101 (2015).

37. Wang, T. et al. Gene Essentiality Profiling Reveals Gene Networks and Synthetic Lethal Interactions with Oncogenic Ras. Cell 168, 890–903.e15 (2017).

38. Hart, T. et al. High-Resolution CRISPR Screens Reveal Fitness Genes and Genotype-Specific Cancer Liabilities. Cell 163, 1515–1526 (2015).

39. Aguirre, A. J. et al. Genomic Copy Number Dictates a Gene-Independent Cell Response to CRISPR/Cas9 Targeting. Cancer Discov. 6, 914–929 (2016).

40. Tzelepis, K. et al. A CRISPR Dropout Screen Identifies Genetic Vulnerabilities and Therapeutic Targets in Acute Myeloid Leukemia. Cell Rep. 17, 1193–1205 (2016).

41. Rauscher, B., Heigwer, F., Breinig, M., Winter, J. & Boutros, M. GenomeCRISPR - a database for high-throughput CRISPR/Cas9 screens. Nucleic Acids Res. 45, D679–D686 (2016).

42. Hart, T. & Moffat, J. BAGEL: a computational framework for identifying essential genes from pooled library screens. BMC Bioinformatics 17, 164 (2016).

43. J. T. L., et al. sva: Surrogate Variable Analysis. (2023) doi:10.18129/B9.bioc.sva.

44. Rahman, M. et al. A method for benchmarking genetic screens reveals a predominant mitochondrial bias. Mol. Syst. Biol. 17, e10013 (2021).

45. Aregger, M. et al. Systematic mapping of genetic interactions for de novo fatty acid synthesis identifies C12orf49 as a regulator of lipid metabolism. Nat. Metab. 2, 499–513 (2020).

46. Acosta-Alvear, D. et al. Paradoxical resistance of multiple myeloma to proteasome inhibitors by decreased levels of 19S proteasomal subunits. eLife 4, e08153 (2015).

