## Supplementary figures and images for "A scalable platform for efficient CRISPR-Cas9 chemical-genetic screens of DNA damage-inducing compounds"

### Supplementary Figure 1

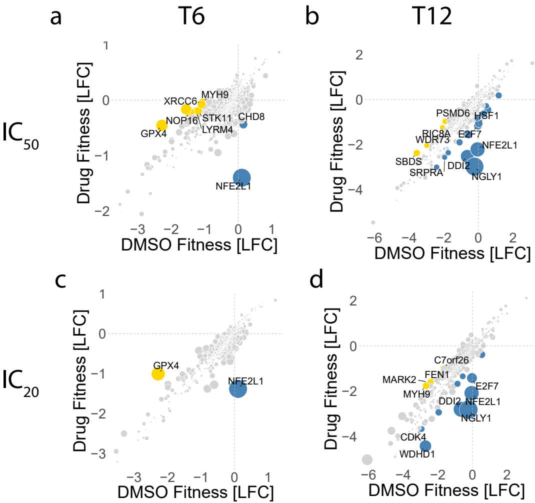

### Supplementary Figure 2

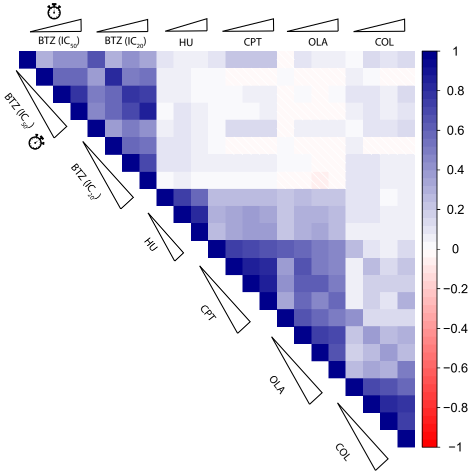

### Supplementary Figure 3

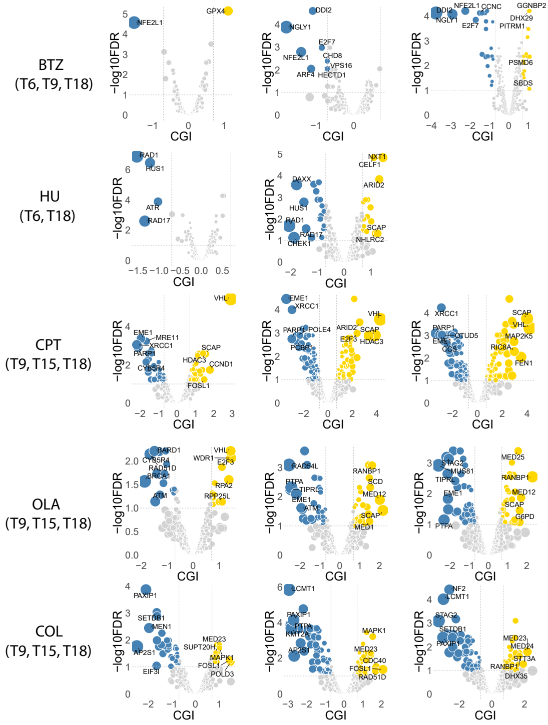

### Supplementary Figure 4

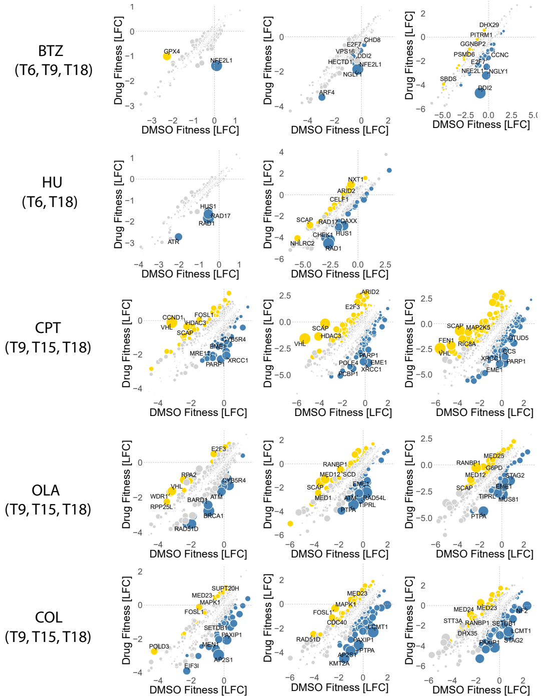

### Supplementary Figure 5

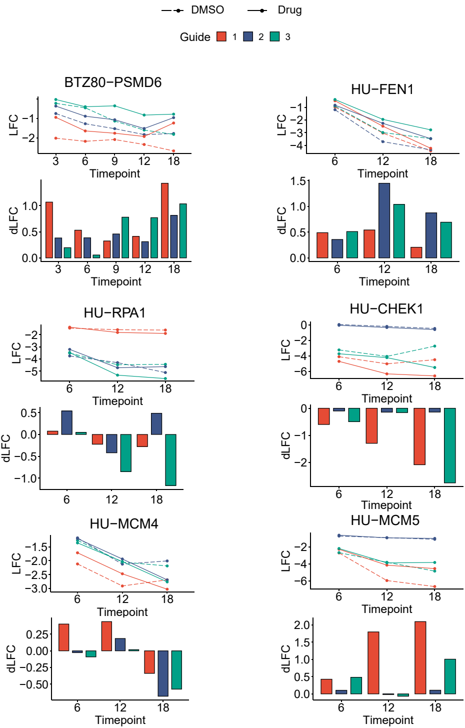

### Supplementary Figure 5

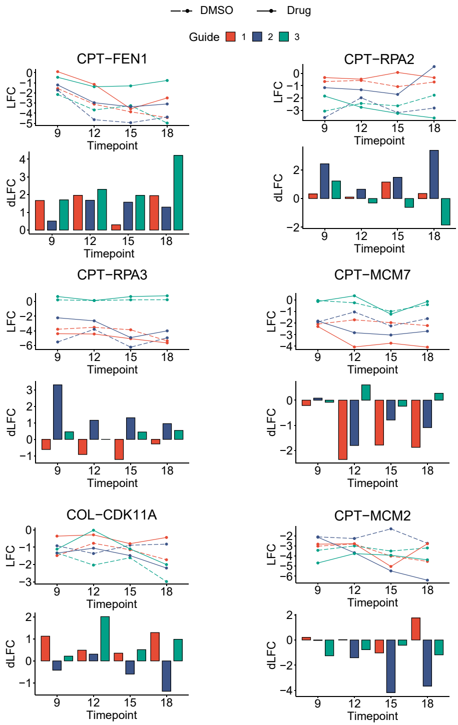

### Supplementary Figure 6

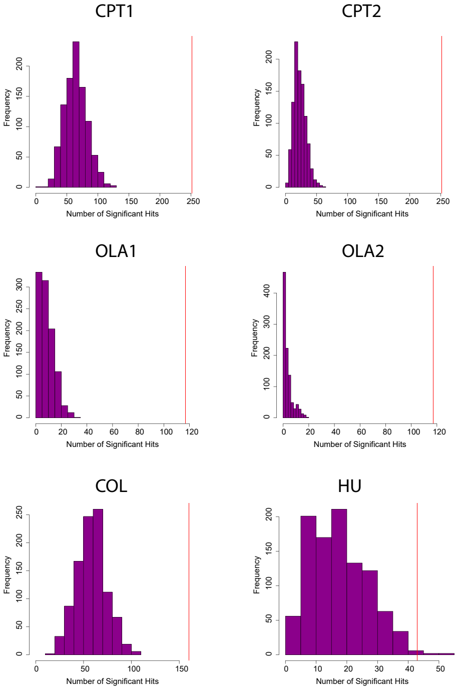

### Supplementary Figure 7

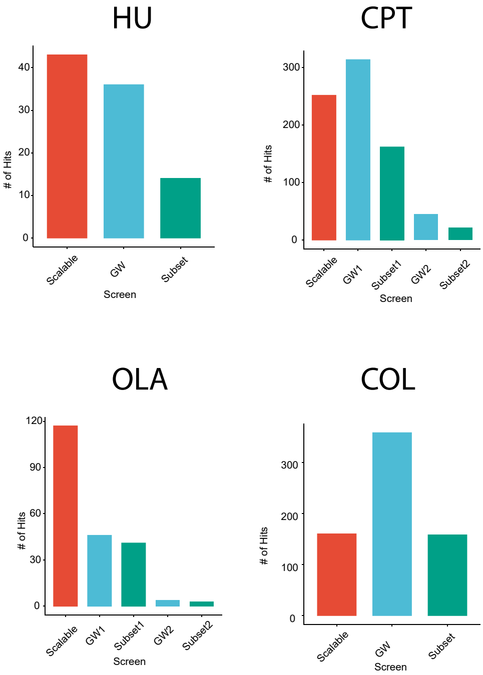

### Supplementary Figure 8

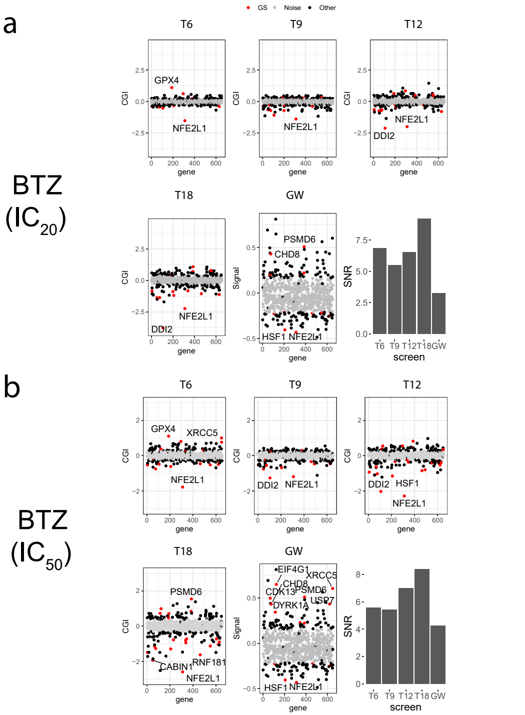

### Supplementary Figure 8

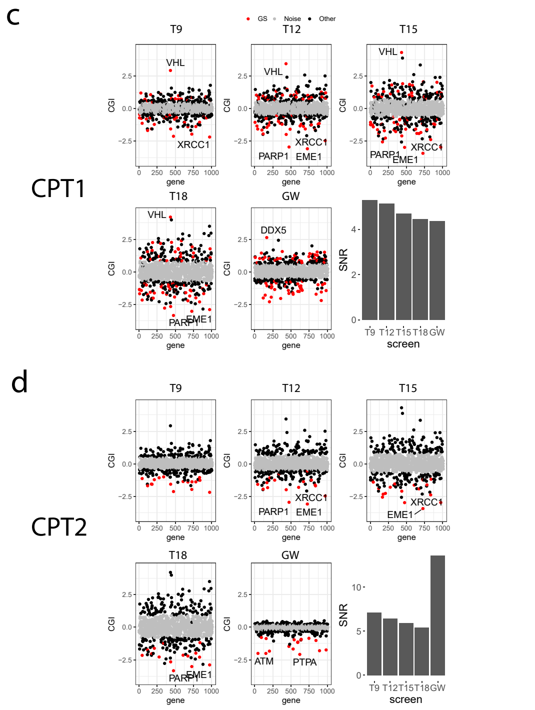

### Supplementary Figure 8

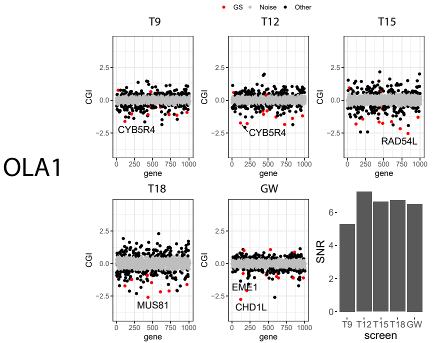

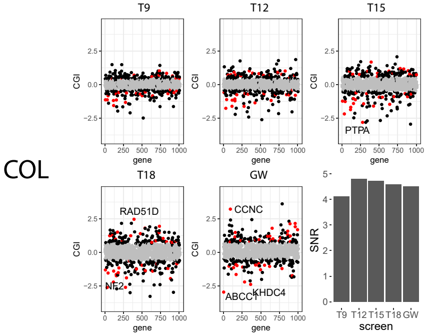

### Supplementary Figure 9

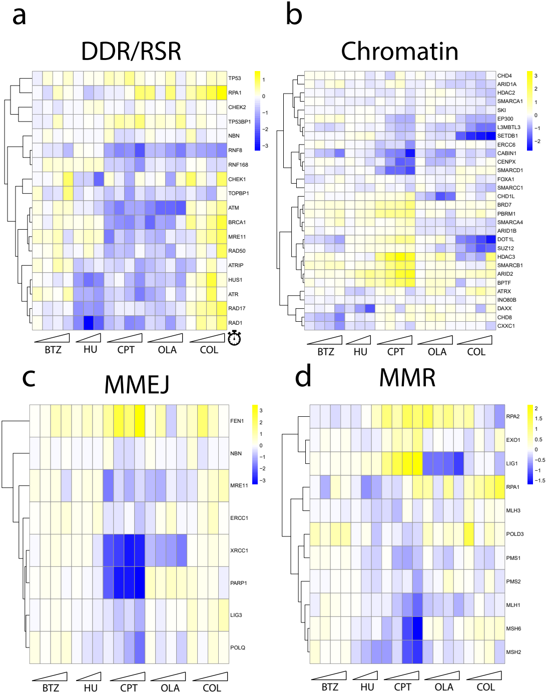

### Supplementary Figure 9

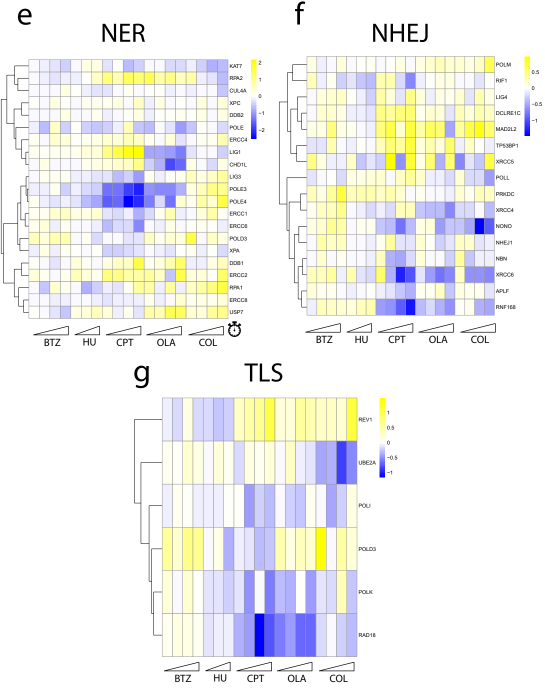
